# Parsing auditory neural code into maximum-entropy packets

**DOI:** 10.1101/2025.11.09.687481

**Authors:** Huanqiu Zhang, Israel Nelken, Tatyana Sharpee

## Abstract

Deciphering the neural code requires identifying its fundamental symbols or code-words. Neural activity is usually interpreted either as a rate code – based on average spike counts – or as a temporal code, which distinguishes patterns with identical counts. Yet, the symbols of the code remain undefined. Here we show that the symbols can be clearly defined by parsing auditory spike trains into variable-duration “packets”, within which precise spike timing is irrelevant. Because packets vary in duration, this is not a rate code. A single neuron could encode very different stimuli depending on the number of spikes it produced per packet. The packet-based code enables real-time readout upon packet completion due to its instantaneous code structure and maximizes information capacity at both single-neuron and population levels.

---

Decades of work have sought to uncover the structure of the neural code. Without a clear answer, studies of encoding and readout depend on assumptions about code structure. The most common assumption is a rate code, where a neuron’s average spike rate is modulated by the stimulus^1,2,3^. An alternative is a temporal code, where spike timing carries information beyond spike counts^4,5,6,7,8,9^. Rate-based analyses often report loss of information compared to the full temporal code conveyed by different spike patterns^10,11,12,13^. Temporal codes can, in principle, capture all information in a spike train, yet they are highly susceptible to spike-time jitter^14^ and complicate decoding and functional analysis. Identifying the structure of the neural code is therefore essential.

The first step in deciphering any code is to identify its basic symbols, or codewords—that is, to determine how sequences are parsed into meaningful units. A classic example is the discovery of codons in protein synthesis: the triplet AUG marks the start of translation, and each subsequent nucleotide triplet specifies a particular amino acid^15^. Efforts to uncover analogous codewords in neural activity initially focused on spike trains from single neurons, yet no consensus emerged on how to segment these sequences. When individual neurons exhibit clear bouts of activity following stimulus onset, it was found that the timing of each bout together with its spike count is sufficient to capture temporal information^13^. At the population level, bouts of activity become far more pronounced with synchronized firing, which has been shown to play a crucial role in temporal encoding and information processing^16,17,18,19,20,21^. Here we show that quiet periods in neural population responses can serve as natural boundaries that delineate codewords, allowing spike trains to be parsed into successive, information-bearing segments.

## Packet organization of neural responses

We simultaneously recorded large neural populations (376 neurons on average per session) from the inferior colliculus, a midbrain hub of the auditory system, using Neuropixels probes^22^ while presenting a diverse set of complex acoustic stimuli, each repeated ten times (Fig. 1a). At the population level, neurons exhibited clear cycles of synchronized activity and silence (Figs. 1b,c; S1a). We segmented population responses into discrete “packets” of activity (Figs. 1c, top; S1b) ^23^. Packets had an average duration of 22*±*8 ms (mean*±*SD, *n* = 1795). Their timing and length were strongly shaped by the incoming sounds: packet onsets were highly reproducible across stimulus repetitions and typically coincided with sharp increases in stimulus energy (Fig. 1b-d). Moreover, packet duration and population spike counts scaled positively with sound energy (Fig. S1c,d). These observations suggest that packet organization may provide a natural parsing of neural activity, offering a first clue to the structure of the neural code.

**Figure 1:**
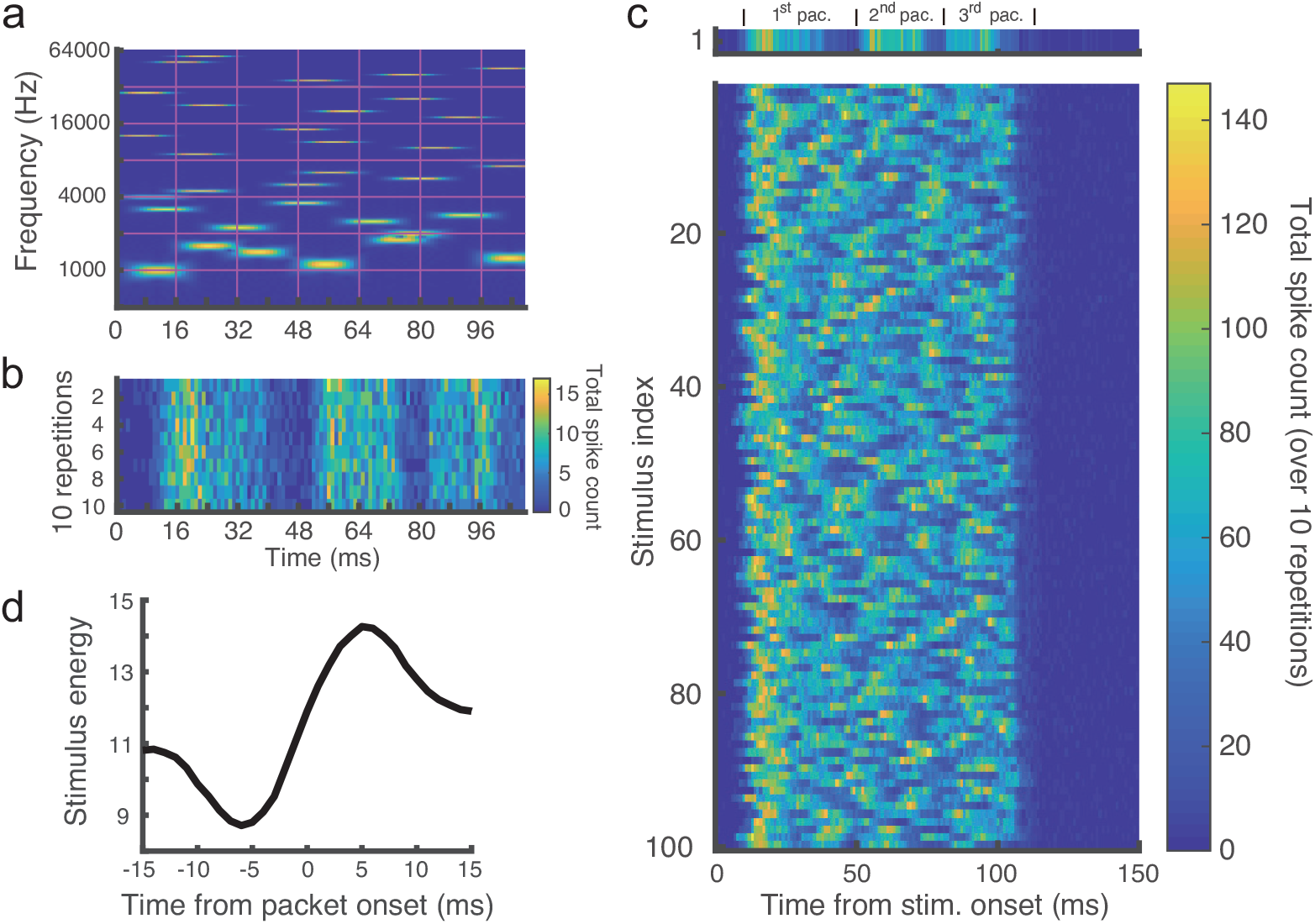
Packet organization of neural population responses. (**a**) Example auditory “tone-cloud” stimulus shown as a spectrogram. (**b**) Summed activity of 508 neurons across 10 repetitions of the stimulus in (**a**), showing highly reproducible packet structure across repeats. (**c**) Population responses (summed over 10 repetitions) to 100 different stimuli (tone clouds). The response to the first stimulus (from **a**) is magnified (top) to illustrate packet definition: the interval from the onset of one high-activity period to the onset of the next. (**d**) Packet-triggered average of stimulus energy, aligned to packet onsets, reveals that packets typically begin ~5 ms after a rise in stimulus energy.

### Packets define an instantaneous neural code

Neural responses within packets have the structure of what communication theory calls an instantaneous code (also known as a prefix-free code)^24^. In such a code, codewords may have different lengths, yet each can be unambiguously identified as soon as it is completed. This requires that no codeword serves as the prefix of another (Fig. 2a).

**Figure 2:**
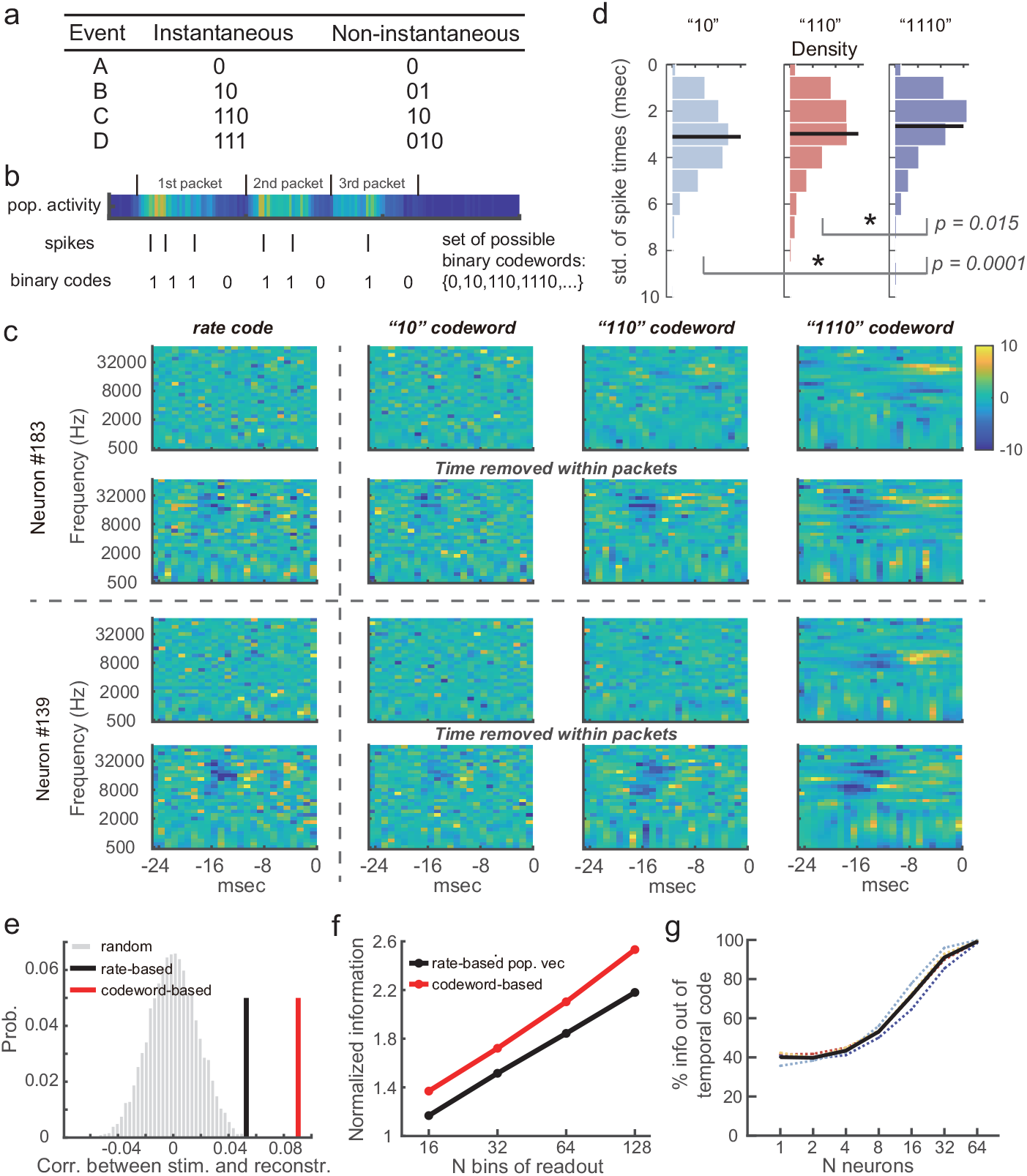
Packet-based neural code captures full information and improves decoding compared to rate code. (**a**) Examples of instantaneous (prefix-free) and non-instantaneous codes. With an instantaneous code, sequences can be read out immediately and unambiguously from a binary string. (**b**) Schematic illustration of interpreting packet-based neural responses as an instantaneous code, based on single-neuron activity aligned to packet structure. (**c**) Spectrotemporal receptive fields (STRFs) for two example neurons, with 1.31 (#183) and 1.46 (#139) spikes per active packet on average, estimated either from firing rate (left) or from individual codewords (remaining columns). Even rows show improvements from removing intra-packet codeword timing. (**d**) Density of spike timing variability across repetitions, averaged for all packets for each neuron. Black lines indicate mean values. “10” vs. “110”: *p* = 0.26 (two-sample *t*-test, *t*-value= 1.13, *df* = 1002), other *p*-values shown in plot (*t*-value= 3.81, *df* = 832 for “10” vs. “1110”; *t*-value= 2.43, *df* = 876 for “110” vs. “1110”). (**e**) Codeword-based stimulus reconstructions (red) were more accurate than rate-based reconstructions (black). (**f**) Information conveyed by population readout vectors based on codewords (red) was higher than rate (black). (**g**) Fraction of temporal-code information preserved by packet-based code as a function of population size. Dashed lines, individual sessions; black line, session average.

To see how this principle arises in neural activity, consider quiescent periods of population activity as “0”, and the number of spikes produced by a neuron within each packet as successive “1”s. This mapping generates the instantaneous code {0, 10, 110, 1110, · · ·} (Fig. 2b), where each trailing “0” marks the end of a codeword – analogous to spaces in written text. Crucially, “0”s are defined at the population level rather than between spikes of a single neuron. If defined at the single-neuron level, fluctuations in spike timing could render the code ambiguous – for instance, an intended “110” could be misread as “1010” if two spikes were separated by a somewhat longer interval. By anchoring “0” to population-wide silence, the code remains robust: variability in individual spike timing does not alter the codeword, as long as all spikes occur within the packet.

### Packet code differs from rate code

This packet-based representation can be summarized by the number of spikes each neuron contributes within a given packet. Yet it differs from a pure rate code in four important ways. First, packets vary in duration, so a neuron may fire at the same average rate in both short and long packets but produce more spikes in the longer one. Second, the interpretation of a neuron’s spikes depends on their alignment with population-wide activity, embedding single-neuron responses within collective dynamics.

There were also two functional consequences of the differences between packet and rate code. The first functional consequence was that the stimulus features associated with different codewords, known as spectrotemporal receptive fields (STRFs), differed among the codewords of the same neuron. These STRFs were also different from the one computed for the firing rate. The STRFs computed from codewords containing multiple spikes exhibited clearer structure (Figs. 2c, S2, S3) than those with fewer spikes or the one based on rate calculations. The effect was particularly pronounced for neurons that often produced more than one spike per packet.

To understand why this might be the case, recall that the packet-based codewords should technically discard all spike timing information within packets. However, the STRF calculation up to now kept the actual time of the codeword occurrence within the packet. It turns out that the variability in the spike timing was larger for codewords containing one or two spikes compared to codewords with three spikes (Fig. 2d). Removing the temporal variability^25^ within packets strongly improved the signal-to-noise ratio in STRF reconstructions (Fig. 2c even rows) and revealed clear differences between stimulus features that are associated with different codewords produced by the same neuron.

These improvements gained by transitioning to codewords and ignoring timing within packets can be quantified in three additional ways. First, one can quantify the accuracy of stimulus reconstructions using the STRF features computed in different ways. We found that reconstructions based on codeword STRFs outperformed rate-based STRFs (Figs. 2e, S4a). Second, one can use tools from information theory^24^ to quantify the amount of mutual information these STRF features captured out of the overall information conveyed by occurrences of the corresponding codeword. We found that STRFs computed without access to precise spike timing accounted for higher percentage of the overall information than STRFs computed with preserved intra-packet timing (Fig. S4b).

Finally, one can examine population-level structure using low-dimensional embedding of neural responses and check how accurately stimuli can be reconstructed in that reduced space. Because packet boundaries were reproducible across repetitions, each stimulus could be subdivided into 3–6 short temporal segments corresponding to individual packets. We first embedded neurons or packet-based codewords based on their response distances. With both firing rate and packet-based codewords, responses were better embedded in a 3D hyperbolic space rather than a 3D Euclidean space (Fig. S5), consistent with prior findings on neural population geometry^26^. In codeword embeddings, codewords from the same neuron tended to occupy similar angular coordinates with larger radii assigned to longer codewords (Fig. S5d,e), similar to their intrinsic tree structure. Such positioning within the embedded hyperbolic space, indicates that codewords from the same neuron provide increasingly refined stimulus representations as codewords with more spikes are engaged. From these embeddings, one can construct population readout vectors by combining vectors from all active codewords or neurons using previously described population vector procedures^1,27^. Readout vectors derived from packet-based codewords conveyed more information about packet identity, and hence stimulus segments, than those based on neuronal firing rates (Fig. 2f; see also Methods and Fig. S4c).

Together, these analyses demonstrate that the instantaneous packet code diverges from traditional rate code in nontrivial ways. Codewords from the same neuron with different spike counts were not mere multiples of a base pattern, but instead transmitted distinct information about sensory input.

### Information loss from discarded timing is recovered at the population level

We now show that even though spike timing within packets is discarded by the instantaneous code, the packet-based code retains nearly all of the information in neural responses once the population size exceeds ~30 neurons. To quantify this, we considered a population of *N* neurons and estimated the information carried by the *N*-dimensional spike count vector about packet identity, and compared it to the information conveyed by the full *N*-dimensional temporal code. For single neurons, ignoring spike timing led to substantial loss: packet-based spike counts accounted for only ~40% of the information in the temporal code, in agreement with previous reports from auditory cortex^13^. By contrast, at the population level, packet-based spike counts captured more than 90% of the temporal-code information when *N ≈* 30 neurons were included (Fig. 2g). Thus, the packet-based code combines the implementation simplicity of a rate-like scheme with the high information fidelity of temporal coding at the population scale.

### The packet-based code has maximum-entropy structure

The packet-based code allows instantaneous readout and straightforward implementation. However whether or not it is efficient, is determined by the frequencies with which codewords are used^24^. For an instantaneous code, efficiency is maximized when the probability of using a codeword (*p*) decays exponentially with its length (*l*): *p ∝ e*^−*l*^ (Eqn. 5.19 in ref.^24^). In spike-packet terms, codeword length equals the number of spikes in a packet plus one. We therefore tested whether spike counts per neuron across packets followed this exponential form.

Indeed, 85.0% of neurons (1280/1505; *p* > 0.05 by *χ*^2^ goodness-of-fit with Benjamini–Hochberg correction; Fig. 3a,b) conformed to an exponential distribution. While probabilities of individual codewords varied across neurons, once normalized by mean firing rate they were well described by a shared exponential law, cf. Fig. 3b. Because exponential is also the maximum entropy distribution given a mean, this result indicates that most neurons communicate with an instantaneous code that both minimizes average spike usage and maximizes encoding capacity^28,29,30^.

**Figure 3:**
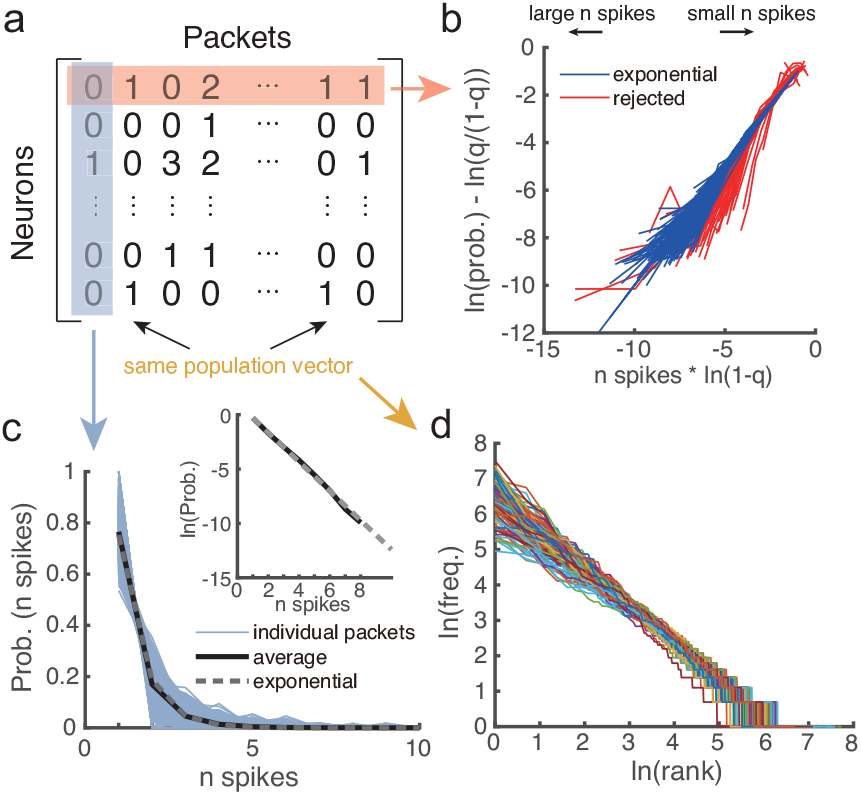
Packet-based code conforms to maximum entropy principle and Zipf’s law. (**a**) Schematic of neural response analysis: each matrix entry is the number of spikes fired by a neuron within a packet. Distributions were examined per neuron (**b**), per packet (**c**), and across packets (**d**). (**b**) Spike counts of individual neurons within active packets followed the exponential distribution predicted by maximum entropy. *χ*^2^ goodness-of-fit tests were performed for each neuron; blue lines, neurons with *p* > 0.05 after correction; red lines, *p* ≤ 0.05. (**c**) Distribution of spike counts from active neurons per packet (colored lines) also conformed to an exponential form. Black line, average distribution across packets; gray dashed line, discrete exponential with the same mean. Insets show log-scale. (**d**) Zipf’s law analysis from 100 iterations, each sampling 12 neurons (different colors). Packets were considered identical if their population spike-count vectors matched exactly. Data shown are from the session with the largest number of simultaneously recorded neurons.

This maximum entropy structure also appeared at the population level. For individual packet IDs, spike counts across active neurons conformed to an exponential distribution in 97.3% of packets (17,457/17,950; Fig. 3c, S6). By contrast, when the same analyses were applied without packet parsing – treating each stimulus response in its entirety – agreement with the exponential law dropped sharply: only 64.3% of neurons (967/1505) and 74.4% of population responses (2976/4000 stimulus repeats) passed the test. Thus, parsing into packets reveals an underlying instantaneous code with maximum entropy statistics that would otherwise be obscured.

At the population-code level, we further observed a power-law relationship between codeword frequency (*F*) and rank (*r*), consistent with Zipf’s law (*F ∝* 1*/r*; Figs. 3d, S6c). This distribution, first described for word frequencies in natural language^31^, has been shown to emerge from maximally informative representations^32^. By analogy, packets may serve as the fundamental “words” of neural communication.

Our findings suggest that the neural code is better understood in terms of codewords rather than individual spikes of neurons. The simplicity of the packet-based codeword structure enables instantaneous readout as soon as each packet is complete, and matches the form for which efficient learning procedures have been developed^14^. Parsing responses based on population-defined “0”s protects against fluctuations in single-neuron spike timing and can be readily implemented by downstream neurons integrating input from hundreds to thousands of presynaptic terminals. Viewing the code in terms of codewords also reframes observations of apparently complex, multi-component neural selectivity^33,34^. What appears as singleneuron integration of multiple stimulus features can instead be parsed into distinct stimulus components associated with different codewords – an approach that improves the recovered information by a simple codeword-based model. Our observations also call for revisions to standard population read-out techniques that assign only one stimulus feature per neuron and weight it by the instantaneous firing rate of the neuron^1^.

Not only should this weighting accounts for noise in neural responses, as was proposed previously^27^, but we now also find that decoding is improved when stimulus-response associations are done for each codeword separately. Finally, although packet duration may vary across brain regions, the ubiquity of packet structure^23^ and oscillatory synchronization across regions and species^35,36,37,38^ suggests that codeword parsing might be a general organizing principle of neural communication.

## Materials and Methods

### Data and preprocessing

We analyzed recordings from a previous study^39^, which provides detailed experimental protocols. Here we summarize the synthesis of tone-cloud stimuli for convenience. Tone clouds were constructed from 36 pure tones, each 16.7 ms long with 5 ms rise/fall times. Frequencies spanned 6 octaves (6 tones/octave). A 100 ms × 6 octave time–frequency region was divided into 36 bins (1 octave × 16.7 ms). Within each octave, tones were randomly permuted and assigned to successive bins, ensuring one tone was active per octave in each bin. Tone onsets were drawn uniformly from 0–16.7 ms to avoid 60 Hz periodicity from synchronous onset. Because onsets varied, tone-cloud duration ranged 104.2–116.4 ms. A pool of 100 tone clouds was generated, stored, and used in all experiments.

For anesthetized recordings (4 sessions from 2 rats), neurons were excluded if their firing rates were not significantly modulated by tone clouds. For each neuron, spike counts were measured during stimulus presentation (150 ms after onset) and a pre-stimulus window of equal length. A paired t-test was performed between the two 1 × 1000 vectors. Only neurons with t-statistics above the 97.5th percentile of the null distribution were retained, yielding 235, 272, 490, and 508 neurons across sessions.

### Packet detection

For each session and stimulus, population firing rates were obtained by summing across neurons and repetitions (Fig. 1c). The time series was smoothed with a 5 ms Hamming kernel, and peaks and troughs were detected. Peaks were required to be *≥*10 ms apart and exceed 5% of the total neuron count. Each peak was assigned to the preceding trough as its onset. The onset of the first packet was defined as the first point after stimulus onset where smoothed activity exceeded 1.5% of neurons. The end of the last packet was the last point within 20 ms after stimulus offset where activity exceeded this threshold.

### Low-rank maximum noise entropy (MNEr) model for estimating STRFs

The MNEr model is described in^40^. Briefly, it is a second-order model of the probability of a binary response given a stimulus:

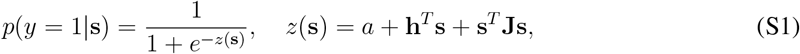

where parameters **a, h, J** are fit by maximizing log-likelihood with **J** constrained to low rank. **h** and **J** capture linear and quadratic receptive fields, respectively (Fig. S7a,b). Quadratic terms contributed little to inferior colliculus (IC) responses and were omitted (Fig. S7c-g).

Stimulus representations **s** were computed via short-time Fourier transform (29 logarithmically spaced frequencies, 500–64,000 Hz; 1 ms windows). At each step, **s** included current and up to 25 ms prior stimulus. All inputs were z-scored.

Neuron nonlinearities were fit with:

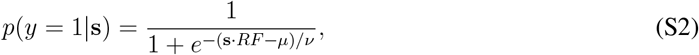

where *RF* is the linear receptive field, and *µ, ν* define neuron-specific nonlinearity.

For rate-based STRF estimation, msec-by-msec responses of a neuron across all stimuli and repetitions were concatenated for the variable *y*. For codeword analyses, spikes were further classified by total spike count per packet. For example, spikes in “10” codewords were retained as events; for “110”, only the average time of the two spikes was treated as the event. To remove time information of events within packets, we computed the overall latency of a neuron’s spikes in all packets, which is used for all events (“10”, “110”, etc.) in their respective packet. Instead of individual codeword (“10”, “110” or “1110”), we also estimated STRFs when events are defined as having “1 or more”, “2 or more”, etc. number of spikes in a packet and obtained qualitatively similar STRFs (Fig. S8).

### Stimulus reconstruction

Reconstructions began with a zero matrix. For spike-train reconstructions, linear receptive fields were shifted to spike times and summed across neurons. For codeword reconstructions, receptive fields estimated for each codeword were shifted to the codeword’s average spike time and summed. Reconstruction accuracy was quantified by Pearson’s correlation between reconstructed and actual stimuli (Gaussian-smoothed, downsampled to 29 frequency bins), averaged across 1000 presentations. Information-preserving recon-structions weighted receptive fields by 1*/ν* from Eqn. S2.

### Pairwise distance matrices

For each neuron, a response vector was defined as spike counts across packets. Pairwise correlations were computed, and distances defined as:

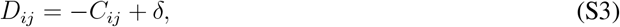

where *C*_*ij*_ is correlation and *δ* ensures positivity.

For codewords, binary response vectors indicated codeword presence/absence per packet. Distances were computed as above. By construction, distinct codewords from the same neuron were anti-correlated; embeddings were thus performed one codeword at a time.

### Hyperbolic non-metric multidimensional scaling (MDS)

Non-metric MDS was used to embed responses in hyperbolic space. Initial positions were obtained via Euclidean MDS (3D, Kruskal’s STRESS1), rescaled to unit radius, and mapped into 3D hyperbolic space (*K* = −1). Distances were computed using the hyperbolic law of cosines:

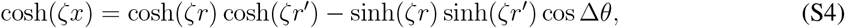

where *x* is hyperbolic distance, *ζ* = 1, *r, r*^*′*^ are radial distances, and Δ*θ* is angular separation.

Coordinates were optimized until rank-order error <0.00001 or 2000 iterations. To prevent over-expansion, an alternating half-fixed optimization was repeated 100 times. Embedding quality was monitored by Pearson correlation between data distances and embedding distances (Fig. S9).

For codeword embeddings, neurons with mostly binary outputs (< 20 multi-spike packets) were embedded first as references. Other codewords were added one by one using only distances to reference neurons. Codewords occurring <10 times were excluded for reliability. Instead of individual codeword (“10”, “110” or “1110”), we also embedded codeword plus defined as having “1 or more”, “2 or more”, etc. number of spikes in a packet and obtained similar results (Fig. S10).

### Population vector decoding

Population readout vectors were constructed by summing embedding locations of active neurons per packet (standard vector^1^), or by additional weighting 1*/ν*, inversely proportional to noise *ν* of each neuron (information-preserving vector^27^). Vectors were discretized into 16–128 bins defined by equidistributed 3D directions^41^. Mutual information was computed as *I*(*R*; *S*) = *H*(*R*) − *H*(*R*|*S*).

For codeword-based decoding, readout vectors were constructed by summing embedding locations of observed codewords (weighted by 1*/ν*). Only *{*10, 110, 1110*}* codewords and reference neurons were included for reliability.

The improvements from using “information-preserving” version of the population code (which takes into variability in neural responses^27^) compared to standard population vector procedure was similar for rate or packet-based codes, cf. Fig. S4c.

### Information-theoretic analyses

Information per event was computed as^12^:

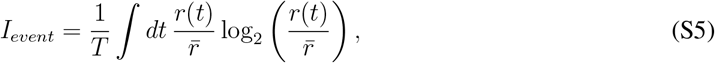

where *T* is stimulus duration, *r*(*t*) is event rate, and 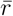 its mean.

Information captured by a linear filter was computed as:

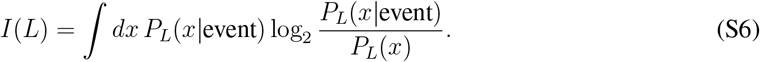

The ratio *I*(*L*)*/I*_*event*_ is used to measure how well the filter captured the actual neuronal responses^42^.

To compare packet-based spike counts with temporal codes, *N* neurons were sampled 100 times. Mutual information was computed for both packet spike counts and temporally binned codes (4 ms bins). Ratios quantified preserved information.

Information estimates were corrected for finite-sample bias by jackknife extrapolation. Subsamples (frac = 80–100% of data) were analyzed and averaged. Results were fit with 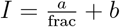, with *b* taken as the infinite-data extrapolation.

### Bayesian fit of discrete exponential distributions

Spike counts per packet were fit with truncated geometric distributions using Bayesian MAP estimation. The likelihood was:

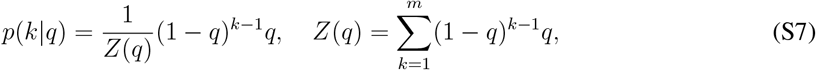

for 1 ≤ *k* ≤ *m*, where *m* is the maximum of observed spike count. Uniform priors were used, and MAP estimates were obtained for *q*.

### Zipf’s law analysis

In each of 100 iterations, 12 neurons were randomly sampled from a session (> 4000 packets). Packets were assigned identities based on identical 12-dimensional spike-count vectors. Identities were ranked by frequency, yielding rank–frequency distributions. Twelve neurons were chosen to balance too few versus too many possible identities.

### Statistics and reproducibility

To test for geometric (or discrete exponential) distribution of spike counts, we used *χ*^2^ goodness-of-fit test, followed by Benjamini–Hochberg correction. To compare information per event captured by STRFs, we used two-sided paired sample t-test with unknown variance. To compare embedding radii of individual codewords in hyperbolic geometry, we used one-sided paired sample t-test with unknown variance. To compare spike timing variability, we used two-sided two-sample t-test with equal but unknown variances. *p*-values associated with Pearson correlation coefficients are all reported two-sided. For each session, neurons were excluded if their firing rates were not significantly modulated by tone clouds (see above). Only codewords below or equal to “1110” were used for population vector decodings.

## Data availability

Auditory data will be made available upon publication in accordance with journal requirements.

## Code availability

Code for low-rank maximum noise entropy model is available at https://github.com/sharpee/low_rank_MNE. Code for hyperbolic MDS is available at https://github.com/sharpee/hyperbolic-MDS. Other codes will be made available on CNL-T github upon publication.

## Acknowledgments

We thank Dr. Robert Gütig for discussions and comments. This research was supported by an AHA-Allen Initiative in Brain Health and Cognitive Impairment award made jointly through the American Heart Association and the Paul G. Allen Frontiers Group (19PABH134610000); the Dorsett Brown Foundation; the Mary K. Chapman Foundation; the Aginsky Fellowship; National Science Foundation (NSF) grant IIS-1724421; the NSF Next Generation Networks for Neuroscience Program (award 2014217); National Institutes of Health grants U19NS112959 and P30AG068635. The funders had no role in study design, data collection and analysis, decision to publish or preparation of the manuscript.

## Author contributions

All authors contributed to the design of this study. I.N. designed the experiments. H.Z. performed the analyses under the supervision of T.S.. All authors together discussed the results and wrote the manuscript.

## Competing interests

The authors declare no competing interests.

## Supplementary figures

**Figure S1:**
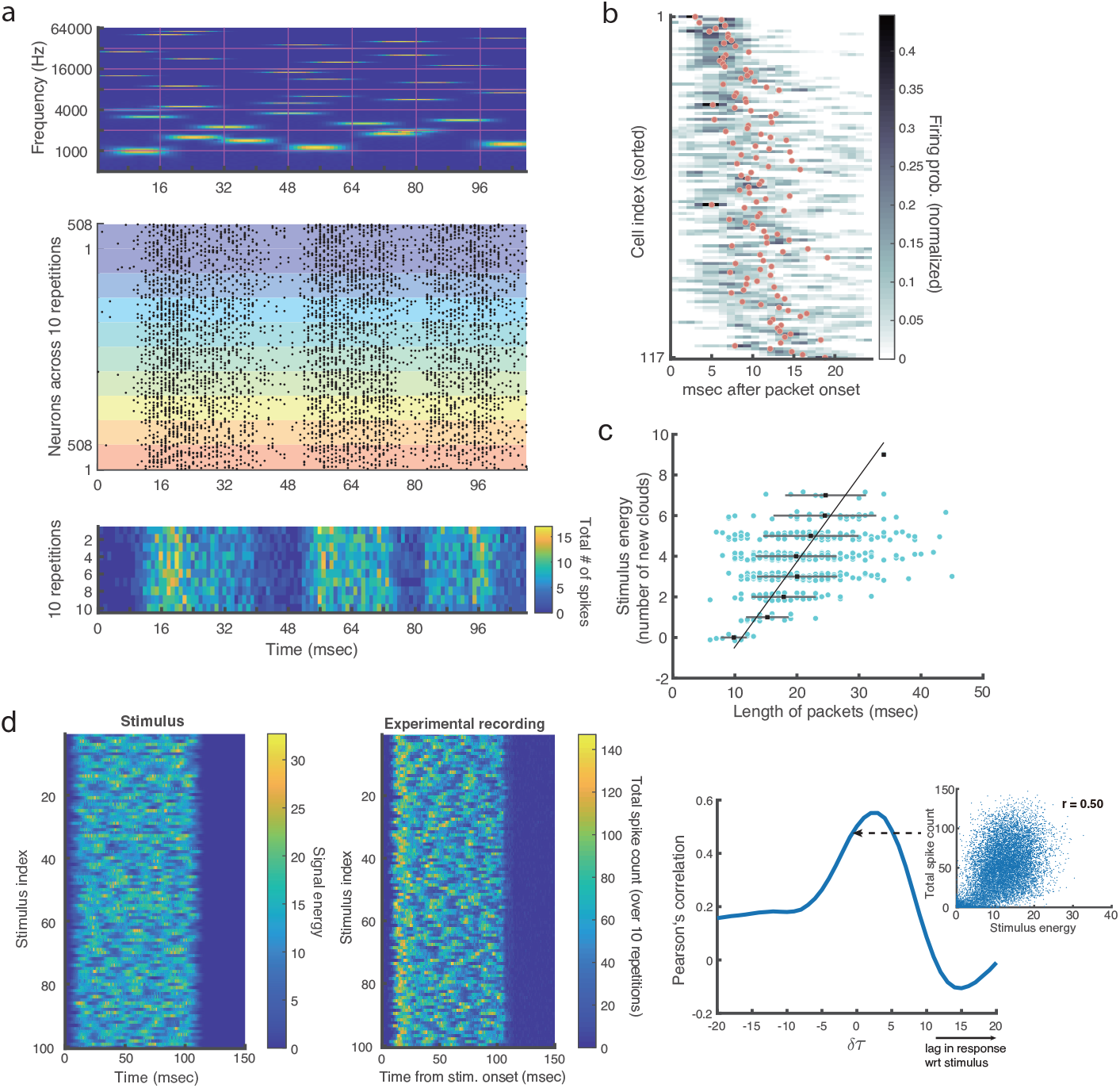
Relationship between packet timing and input statistics. (**a**) Example auditory tone-cloud stimulus (spectrogram, top), raster plot of 508 neurons across 10 repetitions (middle, repetitions indicated by background color), and population firing rate in 1 ms bins (bottom). (**b**) Firing pattern in a representative packet. Shown are 117 neurons firing *≥*3 spikes across 10 repetitions. Grayscale indicates firing probability per ms; red dots mark centroids across repetitions. Neurons are ordered by centroid time across all packets (475 packets × 10 repetitions) to highlight preserved sequential structure^23^. Probabilities are smoothed for visualization. (**c**) Number of new tones starting within *±*6 ms of packet onset versus packet length (*ρ* = 0.34, *p* = 2 *·* 10^−14^). Cyan dots, individual packets (jittered vertically for clarity); black dots and gray lines, mean *±* SD. (**d**) Left: stimulus energy over time for all tone clouds. Middle: population firing rates (same as Fig. 1c). Right: positive correlation between stimulus energy and population spike count. Blue curve shows correlation as a function of stimulus–response delay.

**Figure S2:**
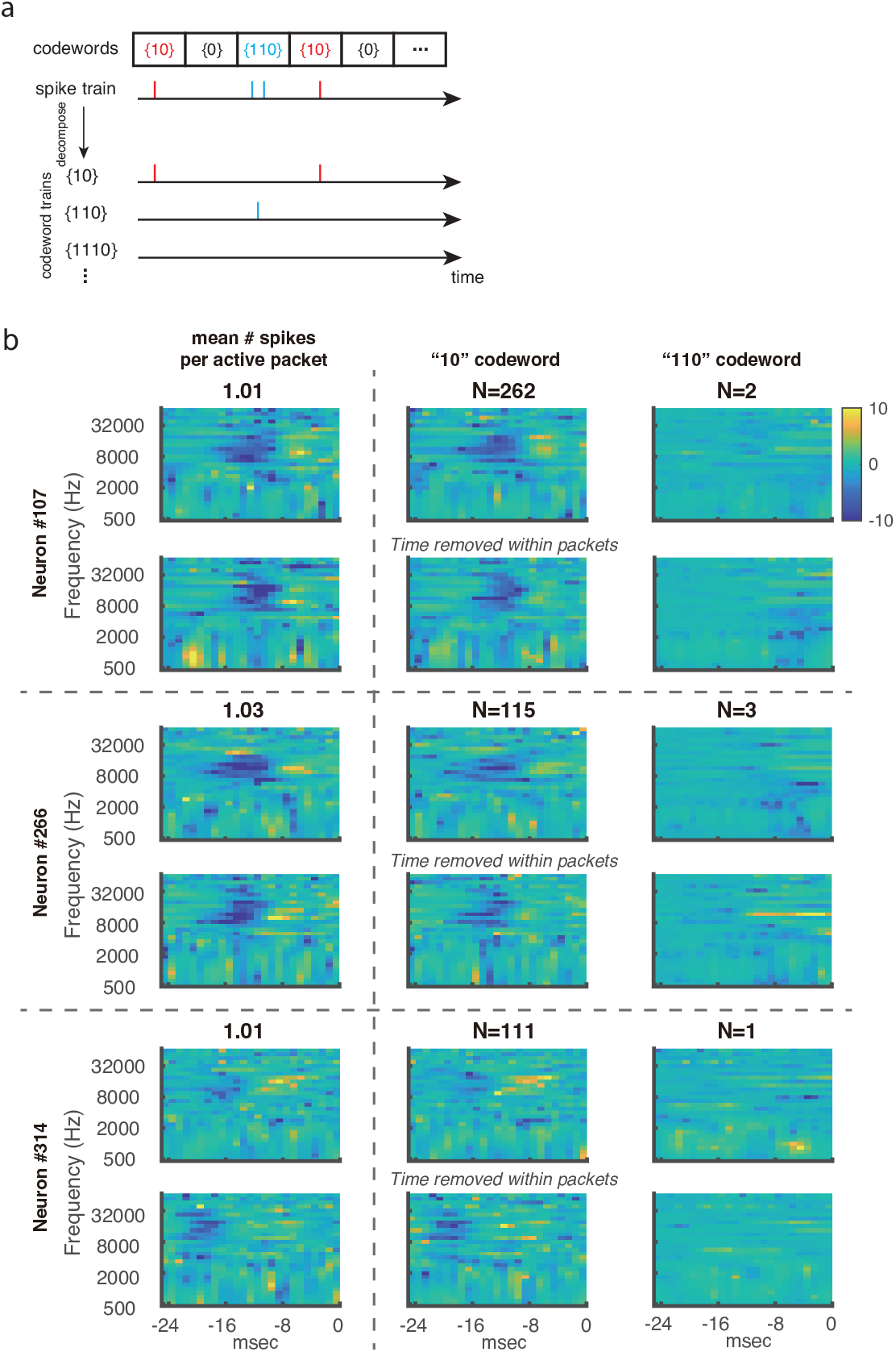
STRFs from different codewords in additional neurons. (**a**) Schematic illustration of decomposing individual neurons’ spike train into subparts corresponding to different codewords to estimate codeword STRFs. (**b**) Spectrotemporal receptive fields (STRFs) for neurons with low activity. Left column: STRFs estimated from full spike trains (activity levels indicated above). Remaining columns: STRFs estimated from “10,” “110,” and “1110” codewords (Methods), with codeword counts shown above.

**Figure S3:**
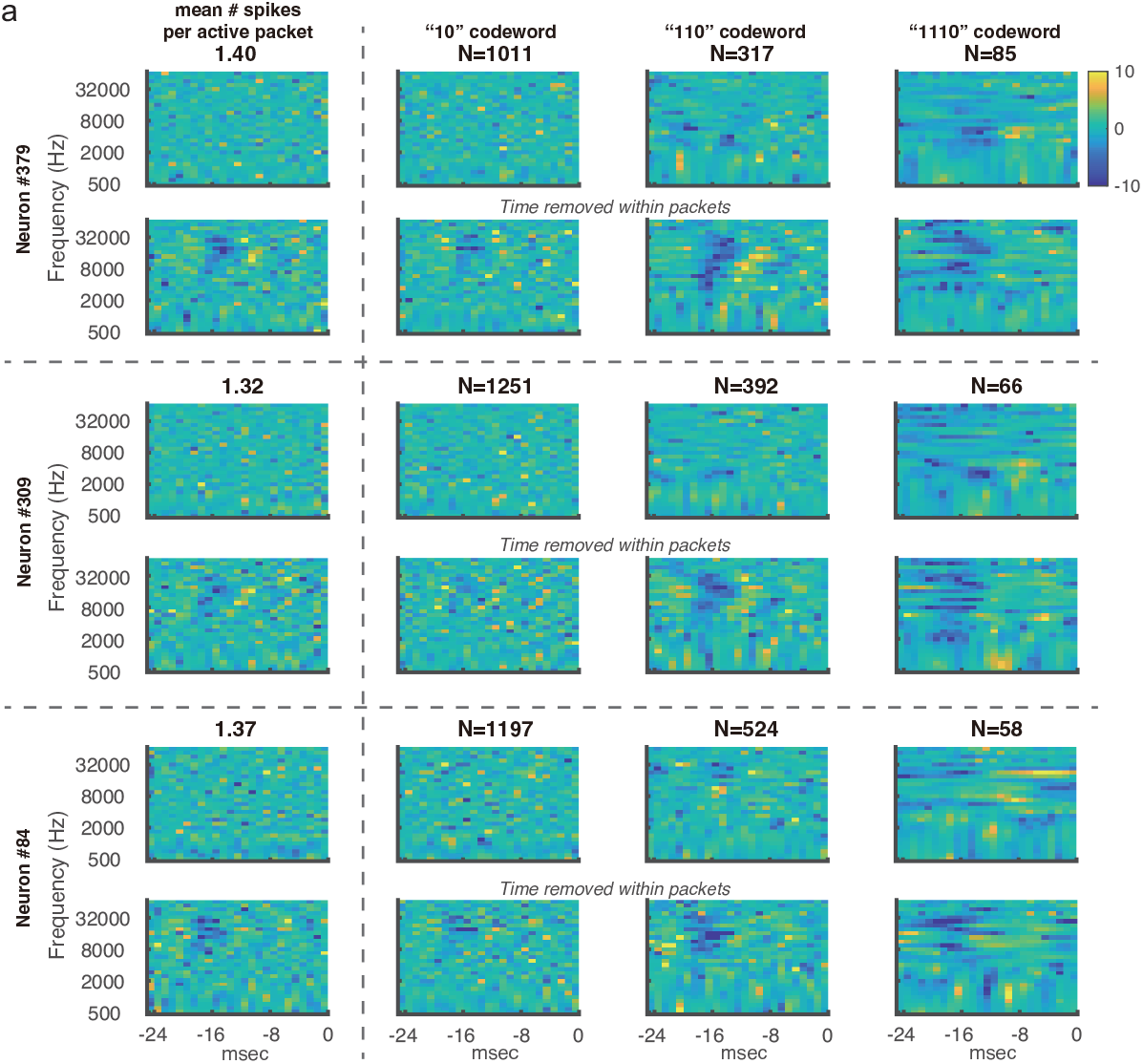
STRFs from different codewords for additional neurons with high activity level. Notations as in Figure S2.

**Figure S4:**
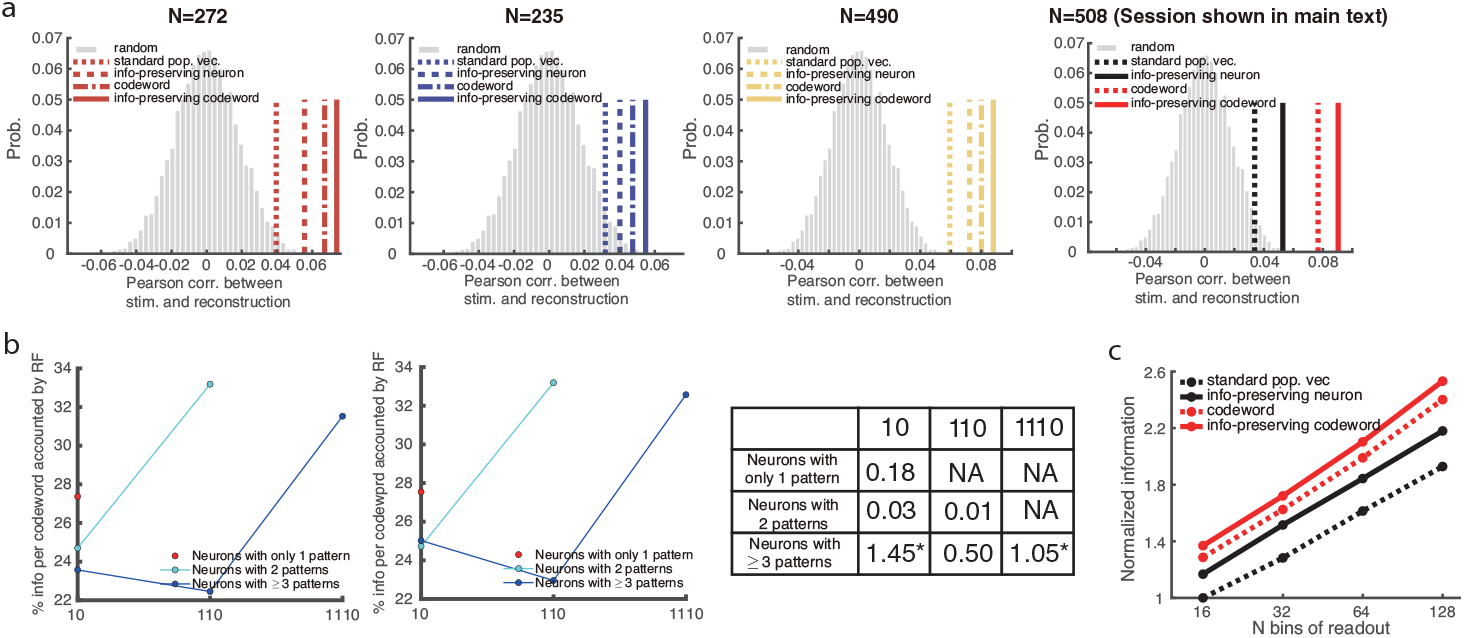
Codewords account for more information than rate coding and improve further when intra-packet timing variation is removed. (**a**) Reconstruction performance across four sessions (neuron counts above). Performance was measured by Pearson’s correlation between reconstructed and actual stimuli. Gray histograms show correlations from 10,000 random vector pairs of matched dimensions. Lines mark mean correlations across stimuli for four linear reconstruction methods (Methods). (**b**) Percentage of information captured by STRFs estimated for individual codewords (Methods). Lines show averages across neurons with the same codeword repertoire. Left: STRFs preserving codeword timing within packets (first row per neuron). Middle: STRFs with codeword timing aligned across packets (second row per neuron). Right: difference between the two; asterisks indicate significant improvement (one-sample t-test). (**c**) Information carried by four linear readouts about packet-defined stimuli based on 3D hyperbolic embeddings, normalized to the information from a standard 16-bin population vector. Two readouts followed an “information-preserving” procedure that accounts for differences in codeword or neuron reliability^27^.

**Figure S5:**
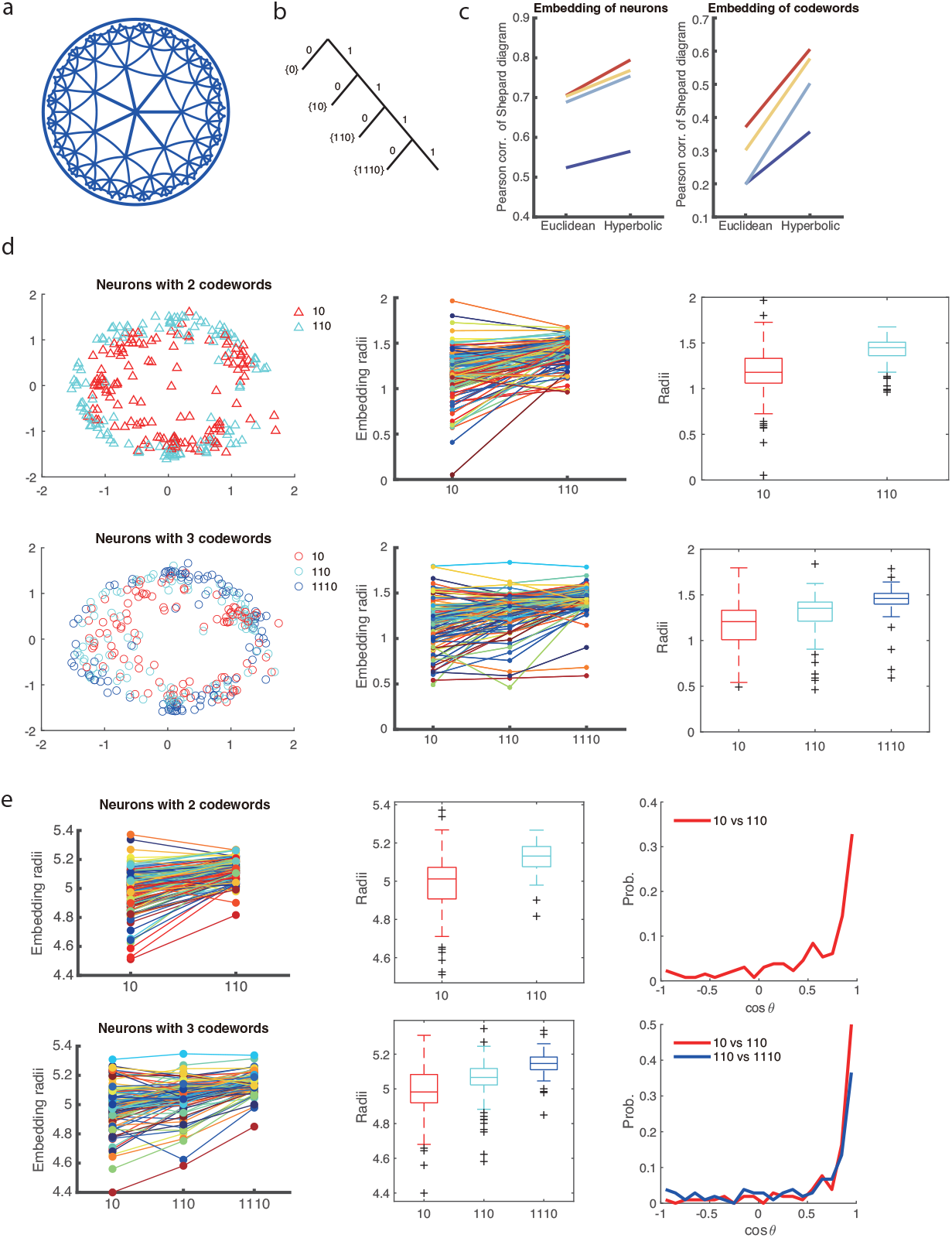
Visualization of the hierarchical structure of codewords using hyperbolic embedding. (**a**) Poincaré disk model of 2D hyperbolic geometry, illustrating its tree-like structure. Curves represent geodesics; all triangles are equilateral and of equal size. (**b**) Schematic of the hierarchical structure underlying the binary instantaneous code. We tested whether longer codewords occupy larger radii in hyperbolic embeddings (see **d**,**e**). (**c**) Shepard diagram correlations for embeddings of neurons (left) and codewords (right). 3D hyperbolic space consistently outperformed 3D Euclidean space. Colors denote different sessions. (**d**) Left: 2D hyperbolic embedding of codewords, grouped by each neuron’s repertoire. Middle: radii of codewords; different codewords from the same neuron are connected by lines of the same color. Significance tests of mean radii: “10” vs. “110” (two-codeword neurons, *p* = 6 *·* 10^−24^, *df* = 130); “10” vs. “110” (three-codeword neurons, *p* = 2 *·* 10^−8^); “110” vs. “1110” (*p* = 7 *·* 10^−16^, *df* = 103). Right: box-plot of radii distributions, with “+” marking outliers. (**e**) Same analyses as (**d**) for 3D hyperbolic embeddings. Significance tests: “10” vs. “110” (two-codeword neurons, *p* = 2 *·* 10^−27^, *df* = 130); “10” vs. “110” (three-codeword neurons, *p* = 4 *·* 10^−10^); “110” vs. “1110” (*p* = 2 *·* 10^−19^, *df* = 103). Right: probability distribution of angular alignment between different codewords from the same neuron.

**Figure S6:**
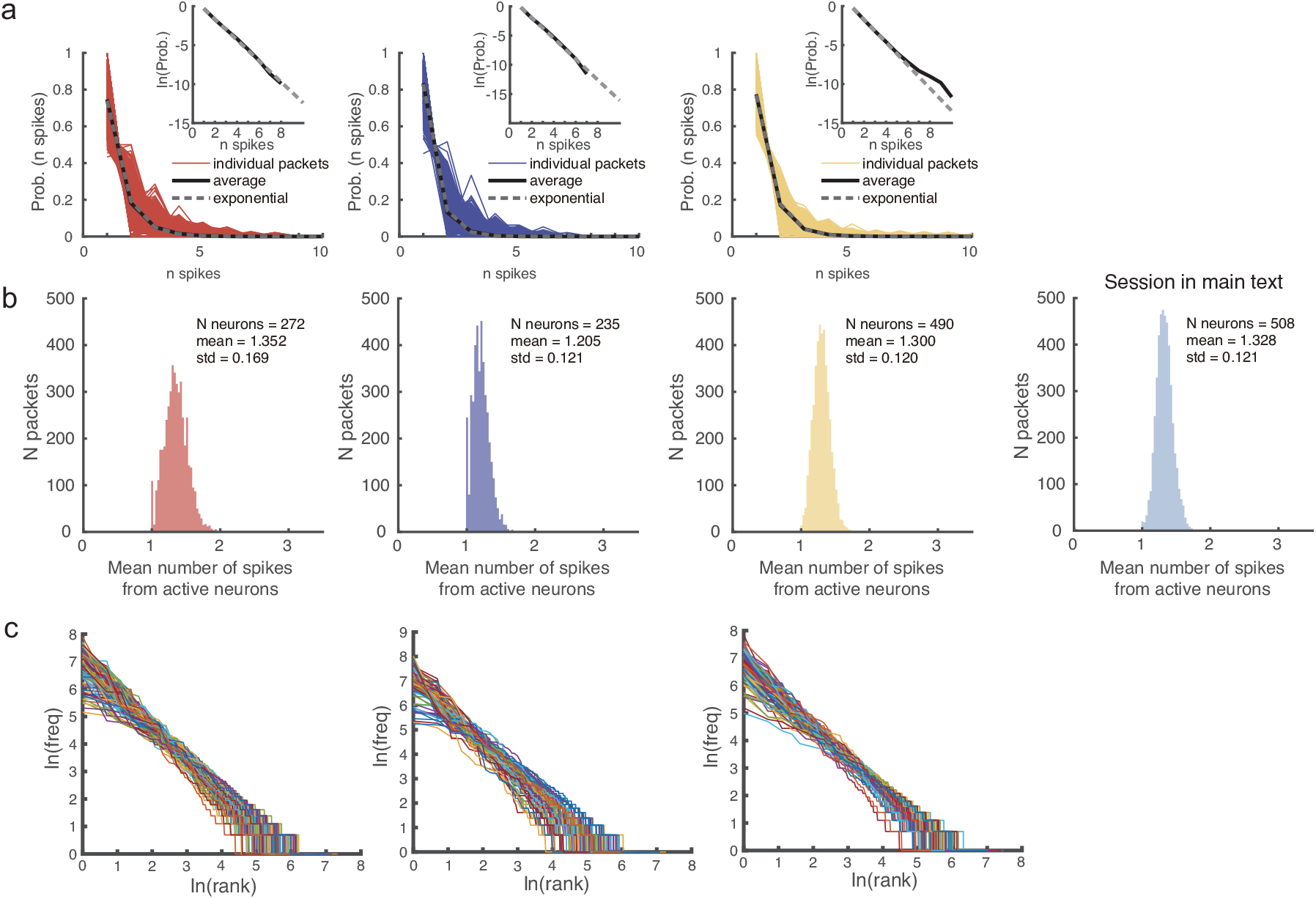
Population maximum entropy distribution, mean activity and Zipfian statistics. (**a**) Distributions of spike counts from active neurons within individual packets for three additional sessions (complementing Fig. 3). (**b**) Histogram of the mean number of spikes fired by active neurons within individual packets. (**c**) Zipf’s law plots for three additional sessions, complementing the example shown in Fig. 3d.

**Figure S7:**
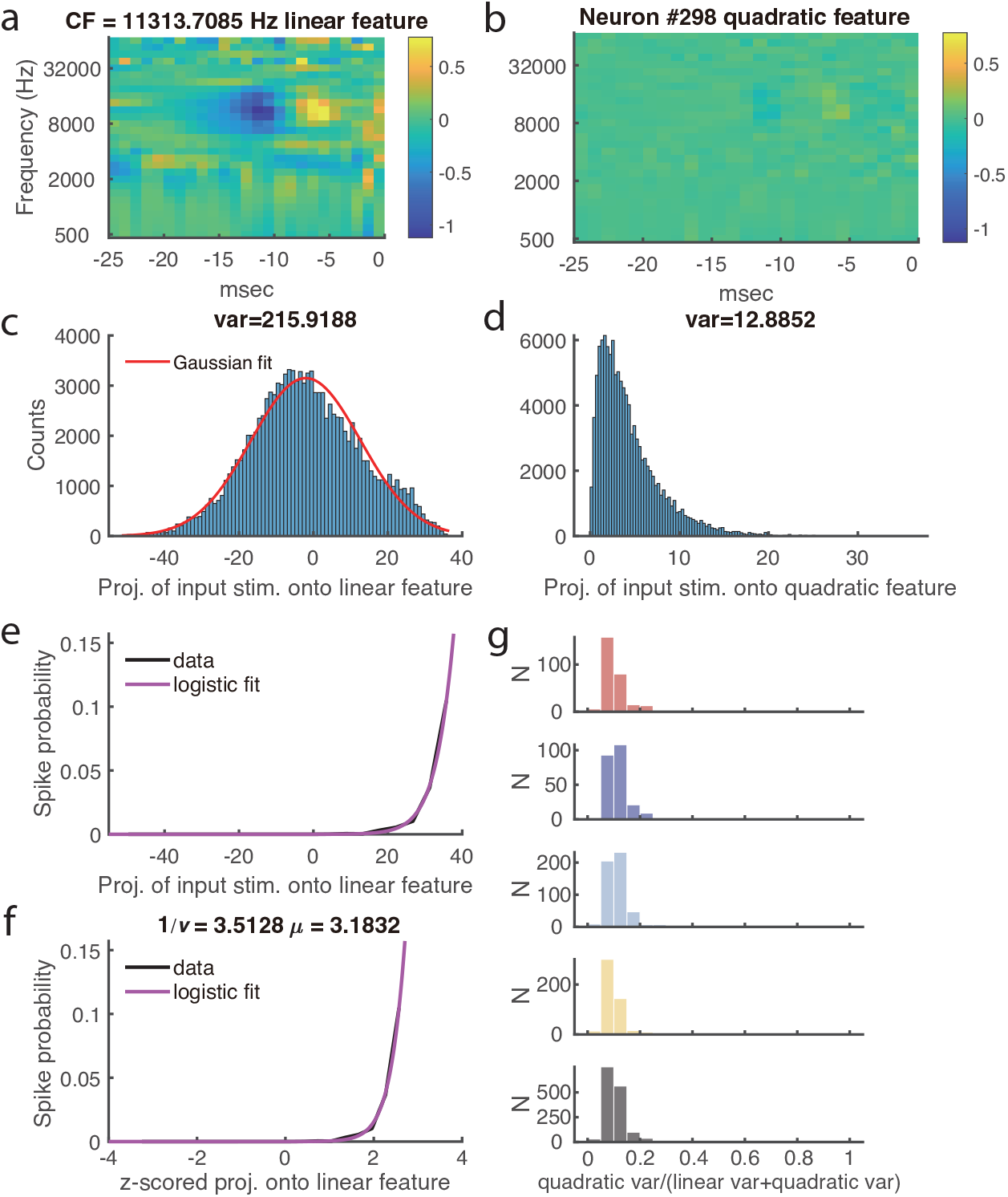
Demonstration of receptive field and nonlinearity analysis with MNEr. (**a**) Linear receptive field of an example neuron estimated with MNEr. Color bar indicates standard deviations from zero. Characteristic frequency was defined as the *y*-coordinate of the maximal pixel after Gaussian smoothing (*σ* = 1 pixel in frequency, 0.75 in time). (**b**) Quadratic receptive field of the same neuron. (**c**) Histogram of projections of tone-cloud stimuli onto the linear receptive field in (**a**). Variance of projections (shown above) quantifies the strength of linear drive. Red line, Gaussian fit. (**d**) Same as (**c**) for the quadratic receptive field in (**b**). Projection variance was much smaller than for the linear field. (**e**) Observed spiking probability as a function of projection onto the linear receptive field (black). Magenta line, logistic fit 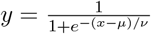, used to estimate nonlinearity parameters *µ* and *ν*. (**f**) Same as (**e**), with *x*-coordinates z-scored to enable comparison of *µ* and *ν* across neurons. (**g**) Contribution of quadratic receptive fields, quantified as the variance ratio of quadratic versus total (linear + quadratic) terms. IC responses were dominated by linear fields, so quadratic terms were omitted from all other analyses. Colors denote sessions; black panel, pooled data.

**Figure S8:**
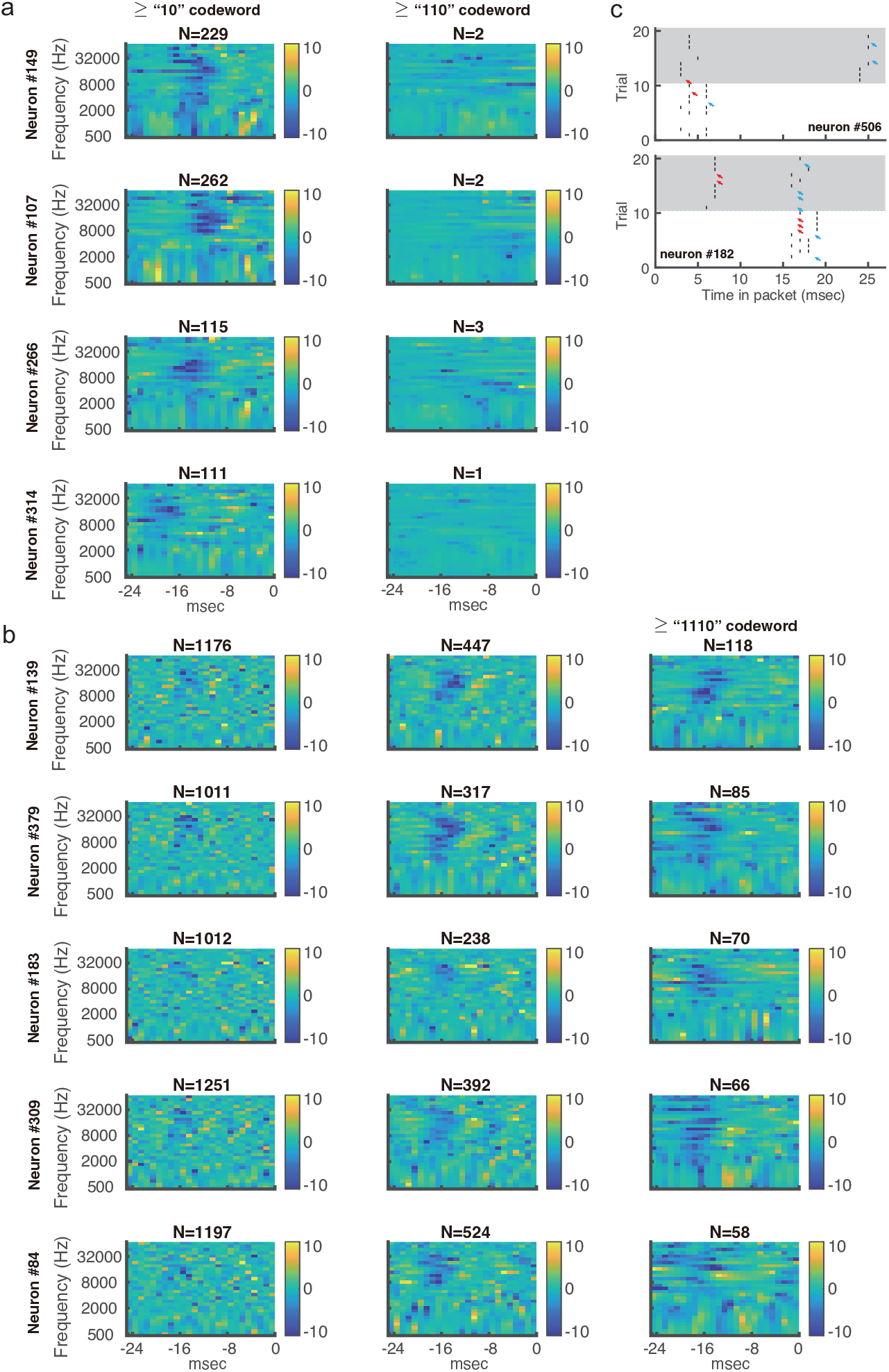
Codewords defined no less than a preset number of spikes. (**a**,**b**) Instead of individual codeword (“10”, “110” or “1110”), codewords are defined here as having “1 or more”, “2 or more”, etc. number of spikes. Spectro-temporal receptive fields (STRFs) estimated for example neurons of low activity level (**a**) or high activity level (**b**) with intra-packet spike timing removed are shown. Other notations are as in Figures S2,S3. (**c**) Raster plots from two example neurons across 10 repetitions of two packets (different background shades). Failures to fire in a repeat converts a “110” codeword into “10” (or even “0”). Red and blue arrows indicate failure of firing the first and second spike, respectively.

**Figure S9:**
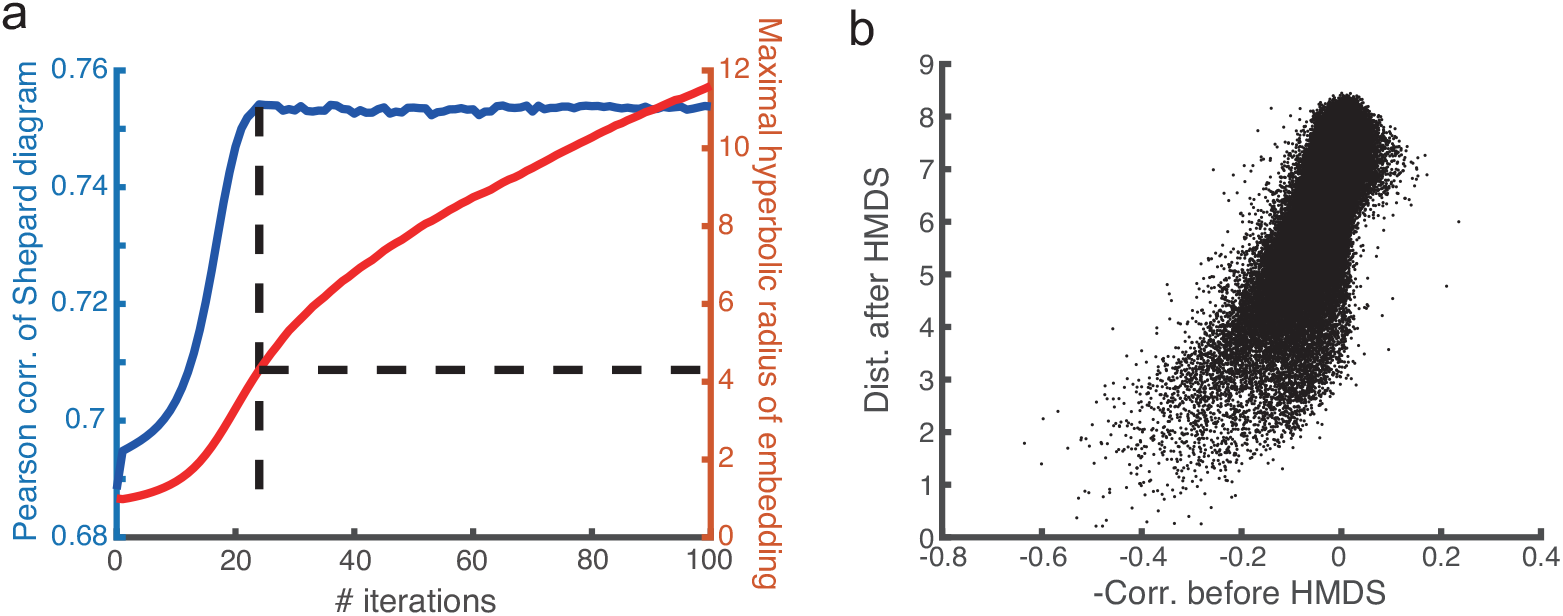
Hyperbolic non-metric MDS. (**a**) Pearson correlation of Shepard diagrams (between distances after iterative non-metric MDS in 3D hyperbolic space and original distances) increased with iteration number, but growth slowed markedly after the vertical dashed line. Iteration 0 corresponds to Euclidean MDS initialization (see Methods). Right *y*-axis: continuous increase of maximal hyperbolic radius beyond the dashed line. (**b**) Shepard diagram at the iteration marked by the dashed line in (**a**).

**Figure S10:**
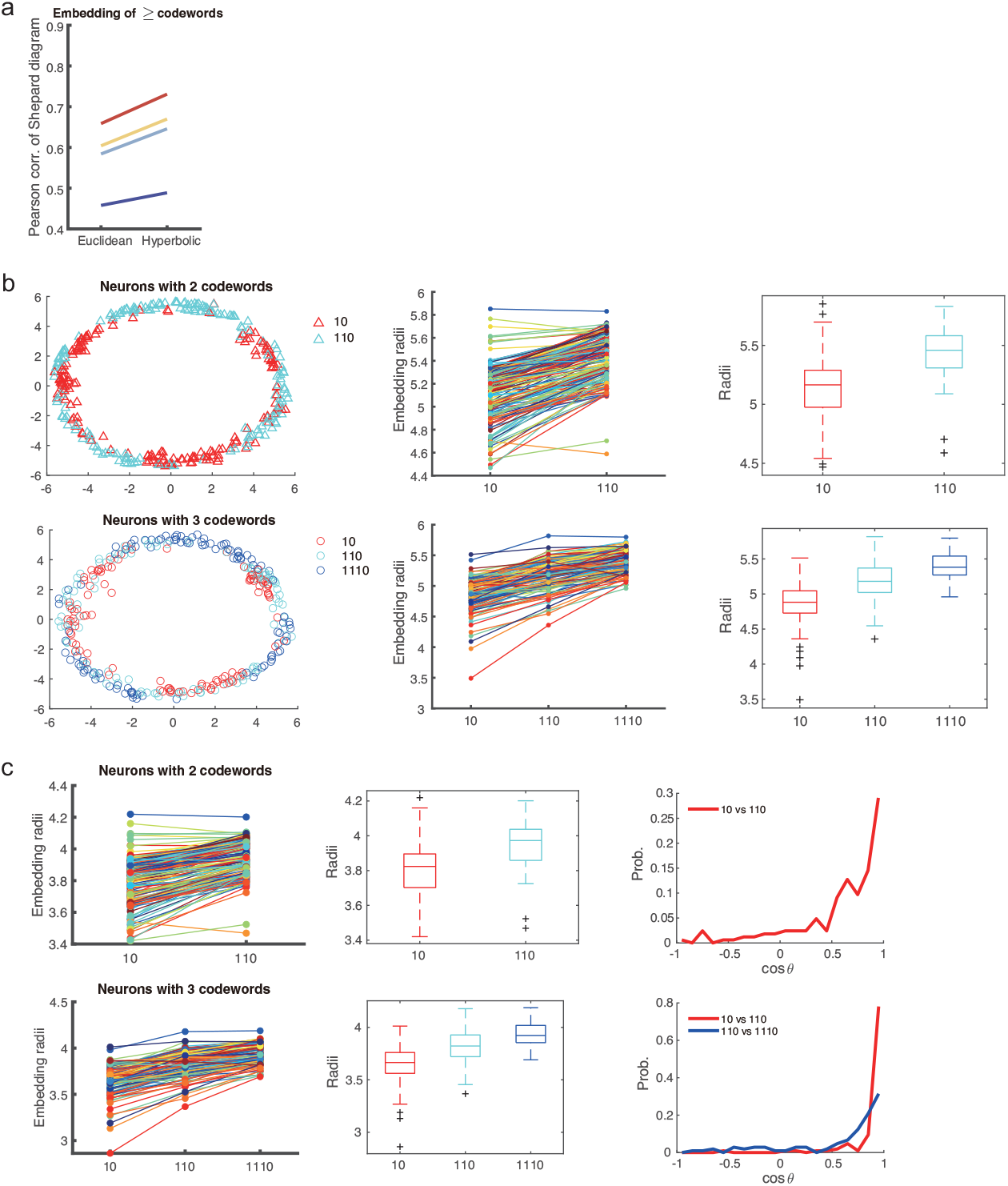
Visualization of the hierarchical structure of codewords plus using hyperbolic embedding. Instead of individual codeword (“10”, “110” or “1110”), codewords are defined here as having “1 or more”, “2 or more”, etc. number of spikes. Notations as in Figure S5. All significance tests of mean radii resulted in *p* < 10^−10^.

## Notes

### Competing Interest Statement

The authors have declared no competing interest.

